## Supplementary material for "Fitness landscape of substrate-adaptive mutations in evolved APC transporters": Figure Supplements

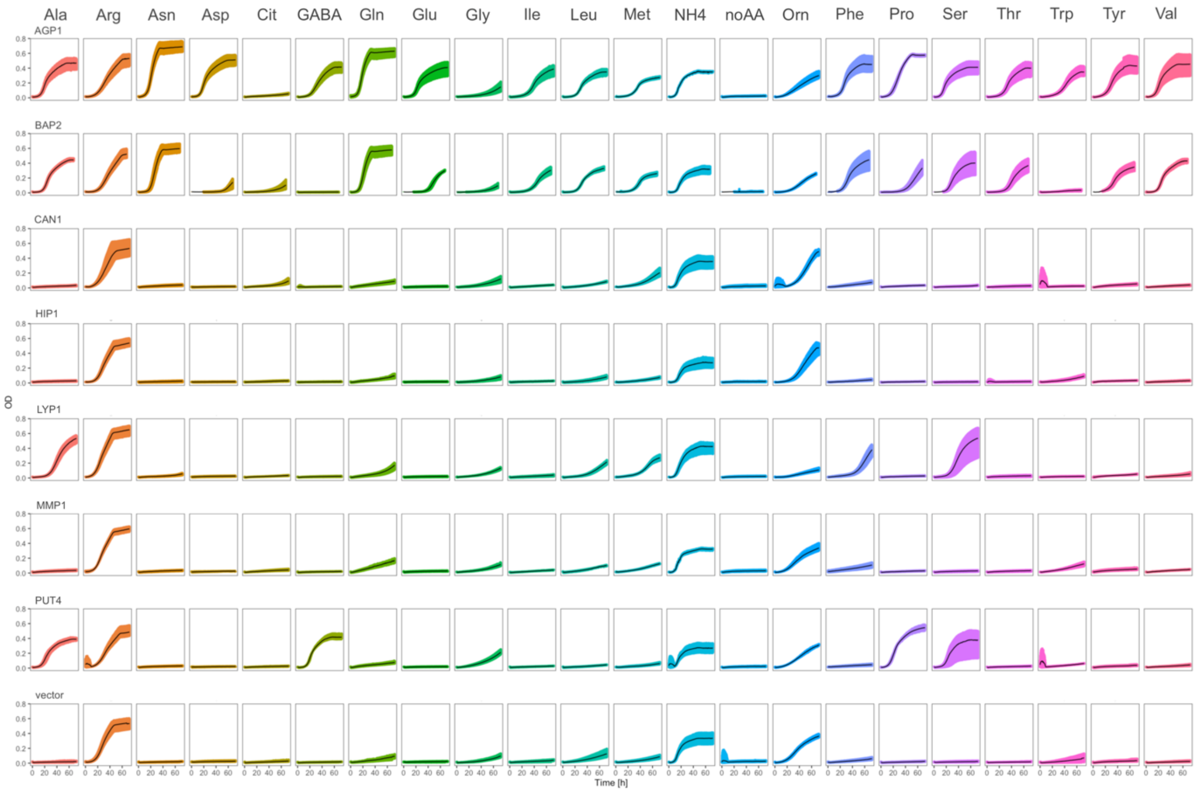


**Figure 1–figure supplement 1**. **YAT transporters support growth on a range of amino acids.** Growth curves of Δ10AA expressing either one of the seven different wild-type yeast amino acid transporter genes (*AGP1*, *BAP2*, *CAN1*, *HIP1*, *LYP1*, *MMP1*, *PUT4*) from pADHXC3GH and the empty vector control on 2 mM of each amino acid. Black lines represent mean values of all measured curves (n ≥ 3). Colored areas represent the SD range.

| | TAT1 | BAP2 | BAP3 | AGP1 | GNP1 | MMP1 | SAM3 | TAT2 | GAP1 | HIP1 | AGP2 | AGP3 | PUT4 | DIP5 | LYP1 | ALP1 | CAN1 |
|---|---|---|---|---|---|---|---|---|---|---|---|---|---|---|---|---|---|
| **TAT1** | 100 | 51 | 49 | 53 | 53 | 36 | 35 | 39 | 43 | 38 | 22 | 27 | 30 | 29 | 31 | 31 | 31 |
| **BAP2** | 51 | 100 | 73 | 53 | 54 | 37 | 38 | 40 | 44 | 43 | 24 | 29 | 31 | 30 | 34 | 33 | 33 |
| **BAP3** | 49 | 73 | 100 | 51 | 53 | 36 | 35 | 39 | 41 | 41 | 22 | 27 | 27 | 29 | 31 | 32 | 30 |
| **AGP1** | 53 | 53 | 51 | 100 | 68 | 39 | 39 | 40 | 44 | 40 | 25 | 27 | 29 | 30 | 32 | 33 | 32 |
| **GNP1** | 53 | 54 | 53 | 68 | 100 | 36 | 36 | 41 | 44 | 39 | 23 | 26 | 27 | 29 | 31 | 32 | 32 |
| **MMP1** | 36 | 37 | 36 | 39 | 36 | 100 | 70 | 38 | 44 | 41 | 24 | 28 | 28 | 31 | 33 | 32 | 32 |
| **SAM3** | 35 | 38 | 35 | 39 | 36 | 70 | 100 | 39 | 47 | 44 | 23 | 27 | 28 | 31 | 32 | 32 | 33 |
| **TAT2** | 39 | 40 | 39 | 40 | 41 | 38 | 39 | 100 | 46 | 43 | 25 | 27 | 30 | 32 | 33 | 35 | 32 |
| **GAP1** | 43 | 44 | 41 | 44 | 44 | 44 | 47 | 46 | 100 | 51 | 28 | 32 | 31 | 32 | 37 | 36 | 36 |
| **HIP1** | 38 | 43 | 41 | 40 | 39 | 41 | 44 | 43 | 51 | 100 | 26 | 29 | 32 | 32 | 33 | 32 | 34 |
| **AGP2** | 22 | 24 | 22 | 25 | 23 | 24 | 23 | 25 | 28 | 26 | 100 | 24 | 31 | 28 | 29 | 28 | 26 |
| **AGP3** | 27 | 29 | 27 | 27 | 26 | 28 | 27 | 27 | 32 | 29 | 24 | 100 | 28 | 31 | 31 | 33 | 33 |
| **PUT4** | 30 | 31 | 27 | 29 | 27 | 28 | 28 | 30 | 31 | 32 | 31 | 28 | 100 | 37 | 34 | 34 | 35 |
| **DIP5** | 29 | 30 | 29 | 30 | 29 | 31 | 31 | 32 | 32 | 32 | 28 | 31 | 37 | 100 | 36 | 36 | 37 |
| **LYP1** | 31 | 34 | 31 | 32 | 31 | 33 | 32 | 33 | 37 | 33 | 29 | 31 | 34 | 36 | 100 | 59 | 62 |
| **ALP1** | 31 | 33 | 32 | 33 | 32 | 32 | 32 | 35 | 36 | 32 | 28 | 33 | 34 | 36 | 59 | 100 | 67 |
| **CAN1** | 31 | 33 | 30 | 32 | 32 | 32 | 33 | 32 | 36 | 34 | 26 | 33 | 35 | 37 | 62 | 67 | 100 |

**Figure 1–figure supplement 2. Pairwise identities of YAT protein sequences from *Saccharomyces cerevisiae*.** The multiple sequence alignment was performed using Clustal Omega, with standard UniProt protein sequences as input (*AGP1*: P25376, *AGP2*: P38090, *AGP3*: P43548, *ALP1*: P38971, *BAP2*: P38084, *BAP3*: P41815, *CAN1*: P04817, *DIP5*: P53388, *GAP1*: P19145, *GNP1*: P48813, *HIP1*: P06775, *LYP1*: P32487, *MMP1*: Q12372, *PUT4*: P15380, *SAM3*: Q08986, *TAT1*: P38085, *TAT2*: P38967). The sequence identity is shown in percentage, from no identity (0) to identical match (100), represented in blue and red respectively.

| N source | Strain | Mean µ | S.E.M. µ |
|---|---|---|---|
| **Ala** | *AGP1* | 0.23 | 0.0048 |
| **Ala** | *BAP2* | 0.25 | 0.0053 |
| **Ala** | *CAN1* | 0.00 | 0.0000 |
| **Ala** | *HIP1* | 0.00 | 0.0000 |
| **Ala** | *LYP1* | 0.15 | 0.0003 |
| **Ala** | *MMP1* | 0.02 | 0.0226 |
| **Ala** | *PUT4* | 0.21 | 0.0068 |
| **Ala** | *vector* | 0.00 | 0.0000 |
| **Asn** | *AGP1* | 0.27 | 0.0043 |
| **Asn** | *BAP2* | 0.29 | 0.0082 |
| **Asn** | *CAN1* | 0.02 | 0.0187 |
| **Asn** | *HIP1* | 0.00 | 0.0000 |
| **Asn** | *LYP1* | 0.03 | 0.0037 |
| **Asn** | *MMP1* | 0.00 | 0.0000 |
| **Asn** | *PUT4* | 0.00 | 0.0000 |
| **Asn** | *vector* | 0.00 | 0.0000 |
| **Asp** | *AGP1* | 0.25 | 0.0022 |
| **Asp** | *BAP2* | 0.15 | 0.0027 |
| **Asp** | *CAN1* | 0.00 | 0.0000 |
| **Asp** | *HIP1* | 0.00 | 0.0000 |
| **Asp** | *LYP1* | 0.00 | 0.0000 |
| **Asp** | *MMP1* | 0.00 | 0.0000 |
| **Asp** | *PUT4* | 0.00 | 0.0000 |
| **Asp** | *vector* | 0.00 | 0.0000 |
| **Cit** | *AGP1* | 0.03 | 0.0044 |
| **Cit** | *BAP2* | 0.07 | 0.0068 |
| **Cit** | *CAN1* | 0.03 | 0.0036 |
| **Cit** | *HIP1* | 0.00 | 0.0000 |
| **Cit** | *LYP1* | 0.00 | 0.0000 |
| **Cit** | *MMP1* | 0.02 | 0.0200 |
| **Cit** | *PUT4* | 0.00 | 0.0000 |
| **Cit** | *vector* | 0.01 | 0.0069 |
| **GABA** | *AGP1* | 0.17 | 0.0062 |
| **GABA** | *BAP2* | 0.00 | 0.0000 |
| **GABA** | *CAN1* | 0.00 | 0.0000 |
| **GABA** | *HIP1* | 0.00 | 0.0000 |
| **GABA** | *LYP1* | 0.00 | 0.0000 |
| **GABA** | *MMP1* | 0.00 | 0.0000 |
| **GABA** | *PUT4* | 0.28 | 0.0155 |
| **GABA** | *vector* | 0.00 | 0.0000 |
| **Gln** | *AGP1* | 0.31 | 0.0194 |
| **Gln** | *BAP2* | 0.31 | 0.0033 |
| **Gln** | *CAN1* | 0.07 | 0.0087 |
| **Gln** | *HIP1* | 0.05 | 0.0094 |
| **Gln** | *LYP1* | 0.04 | 0.0022 |
| **Gln** | *MMP1* | 0.08 | 0.0085 |
| **Gln** | *PUT4* | 0.07 | 0.0078 |
| **Gln** | *vector* | 0.05 | 0.0042 |
| **Glu** | *AGP1* | 0.23 | 0.0034 |
| **Glu** | *BAP2* | 0.21 | 0.0070 |
| **Glu** | *CAN1* | 0.00 | 0.0000 |
| **Glu** | *HIP1* | 0.00 | 0.0000 |
| **Glu** | *LYP1* | 0.00 | 0.0000 |
| **Glu** | *MMP1* | 0.00 | 0.0000 |
| **Glu** | *PUT4* | 0.00 | 0.0000 |
| **Glu** | *vector* | 0.00 | 0.0000 |
| **Gly** | *AGP1* | 0.05 | 0.0033 |
| **Gly** | *BAP2* | 0.07 | 0.0045 |
| **Gly** | *CAN1* | 0.05 | 0.0020 |
| **Gly** | *HIP1* | 0.04 | 0.0008 |
| **Gly** | *LYP1* | 0.04 | 0.0041 |
| **Gly** | *MMP1* | 0.04 | 0.0029 |
| **Gly** | *PUT4* | 0.06 | 0.0047 |
| **Gly** | *vector* | 0.05 | 0.0026 |
| **Ile** | *AGP1* | 0.14 | 0.0029 |
| **Ile** | *BAP2* | 0.14 | 0.0019 |
| **Ile** | *CAN1* | 0.02 | 0.0159 |
| **Ile** | *HIP1* | 0.00 | 0.0000 |
| **Ile** | *LYP1* | 0.03 | 0.0143 |
| **Ile** | *MMP1* | 0.02 | 0.0166 |
| **Ile** | *PUT4* | 0.00 | 0.0000 |
| **Ile** | *vector* | 0.01 | 0.0088 |
| **Leu** | *AGP1* | 0.20 | 0.0043 |
| **Leu** | *BAP2* | 0.21 | 0.0036 |
| **Leu** | *CAN1* | 0.06 | 0.0091 |
| **Leu** | *HIP1* | 0.05 | 0.0044 |
| **Leu** | *LYP1* | 0.06 | 0.0074 |
| **Leu** | *MMP1* | 0.05 | 0.0066 |
| **Leu** | *PUT4* | 0.04 | 0.0045 |
| **Leu** | *vector* | 0.05 | 0.0050 |
| **Met** | *AGP1* | 0.17 | 0.0031 |
| **Met** | *BAP2* | 0.19 | 0.0050 |
| **Met** | *CAN1* | 0.06 | 0.0043 |
| **Met** | *HIP1* | 0.05 | 0.0017 |
| **Met** | *LYP1* | 0.08 | 0.0020 |
| **Met** | *MMP1* | 0.05 | 0.0034 |
| **Met** | *PUT4* | 0.03 | 0.0010 |
| **Met** | *vector* | 0.05 | 0.0025 |
| **Phe** | *AGP1* | 0.21 | 0.0026 |
| **Phe** | *BAP2* | 0.18 | 0.0023 |
| **Phe** | *CAN1* | 0.05 | 0.0043 |
| **Phe** | *HIP1* | 0.02 | 0.0121 |
| **Phe** | *LYP1* | 0.08 | 0.0025 |
| **Phe** | *MMP1* | 0.04 | 0.0056 |
| **Phe** | *PUT4* | 0.03 | 0.0142 |
| **Phe** | *vector* | 0.04 | 0.0041 |
| **Pro** | *AGP1* | 0.19 | 0.0055 |
| **Pro** | *BAP2* | 0.13 | 0.0050 |
| **Pro** | *CAN1* | 0.00 | 0.0000 |
| **Pro** | *HIP1* | 0.00 | 0.0000 |
| **Pro** | *LYP1* | 0.00 | 0.0000 |
| **Pro** | *MMP1* | 0.00 | 0.0000 |
| **Pro** | *PUT4* | 0.22 | 0.0117 |
| **Pro** | *vector* | 0.00 | 0.0000 |
| **Ser** | *AGP1* | 0.27 | 0.0032 |
| **Ser** | *BAP2* | 0.21 | 0.0032 |
| **Ser** | *CAN1* | 0.02 | 0.0056 |
| **Ser** | *HIP1* | 0.00 | 0.0000 |
| **Ser** | *LYP1* | 0.16 | 0.0081 |
| **Ser** | *MMP1* | 0.00 | 0.0000 |
| **Ser** | *PUT4* | 0.21 | 0.0264 |
| **Ser** | *vector* | 0.00 | 0.0000 |
| **Thr** | *AGP1* | 0.17 | 0.0019 |
| **Thr** | *BAP2* | 0.17 | 0.0040 |
| **Thr** | *CAN1* | 0.00 | 0.0000 |
| **Thr** | *HIP1* | 0.02 | 0.0169 |
| **Thr** | *LYP1* | 0.00 | 0.0000 |
| **Thr** | *MMP1* | 0.00 | 0.0000 |
| **Thr** | *PUT4* | 0.00 | 0.0000 |
| **Thr** | *vector* | 0.00 | 0.0000 |
| **Trp** | *AGP1* | 0.11 | 0.0030 |
| **Trp** | *BAP2* | 0.00 | 0.0000 |
| **Trp** | *CAN1* | 0.00 | 0.0000 |
| **Trp** | *HIP1* | 0.04 | 0.0034 |
| **Trp** | *LYP1* | 0.00 | 0.0000 |
| **Trp** | *MMP1* | 0.05 | 0.0053 |
| **Trp** | *PUT4* | 0.08 | 0.0376 |
| **Trp** | *vector* | 0.04 | 0.0144 |
| **Tyr** | *AGP1* | 0.16 | 0.0069 |
| **Tyr** | *BAP2* | 0.16 | 0.0058 |
| **Tyr** | *CAN1* | 0.06 | 0.0285 |
| **Tyr** | *HIP1* | 0.00 | 0.0000 |
| **Tyr** | *LYP1* | 0.04 | 0.0238 |
| **Tyr** | *MMP1* | 0.02 | 0.0201 |
| **Tyr** | *PUT4* | 0.02 | 0.0188 |
| **Tyr** | *vector* | 0.01 | 0.0126 |
| **Val** | *AGP1* | 0.23 | 0.0129 |
| **Val** | *BAP2* | 0.25 | 0.0131 |
| **Val** | *CAN1* | 0.02 | 0.0204 |
| **Val** | *HIP1* | 0.00 | 0.0000 |
| **Val** | *LYP1* | 0.03 | 0.0058 |
| **Val** | *MMP1* | 0.05 | 0.0049 |
| **Val** | *PUT4* | 0.03 | 0.0069 |
| **Val** | *vector* | 0.00 | 0.0000 |

**Figure 1–Source data 1. Growth rates of YAT expressing yeast.** Growth rate values of Δ10AA expressing either one of the seven different wild-type yeast amino acid transporter genes (*AGP1*, *BAP2*, *CAN1*, *HIP1*, *LYP1*, *MMP1*, *PUT4*) from pADHXC3GH and the empty vector control on 2 mM of each amino acid. The growth rates were calculated based on Figure 1-Figure supplement 1.

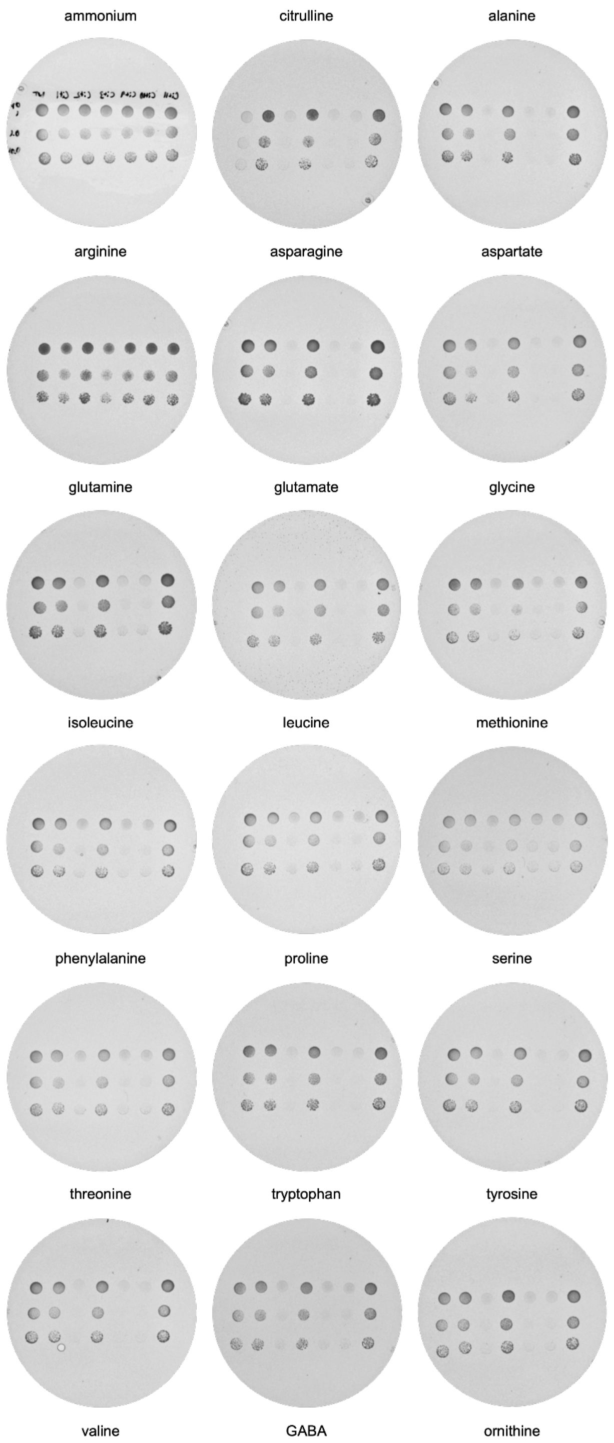


**Figure 2–figure supplement 1**. ***In vivo* evolution of *AGP1*.** Growth assays of Δ10AA pADHXC3GH-*AGP1* variants isolated from Cit evolution, spotted on minimal agar with 1 mM of the respective amino acid as the sole nitrogen source. Dishes were imaged after 6 days of incubation at 30 °C. Order of *AGP1* variants in each dish from left to right: *AGP1*-wild type,

*AGP1*-Cit1, *AGP1*-Cit2, *AGP1*-Cit3, *AGP1*-Cit9, *AGP1*-Cit10, *AGP1*-Cit11. Spotted are 5 μL of $OD_{600}$ of 1, 0.1, and 0.01.

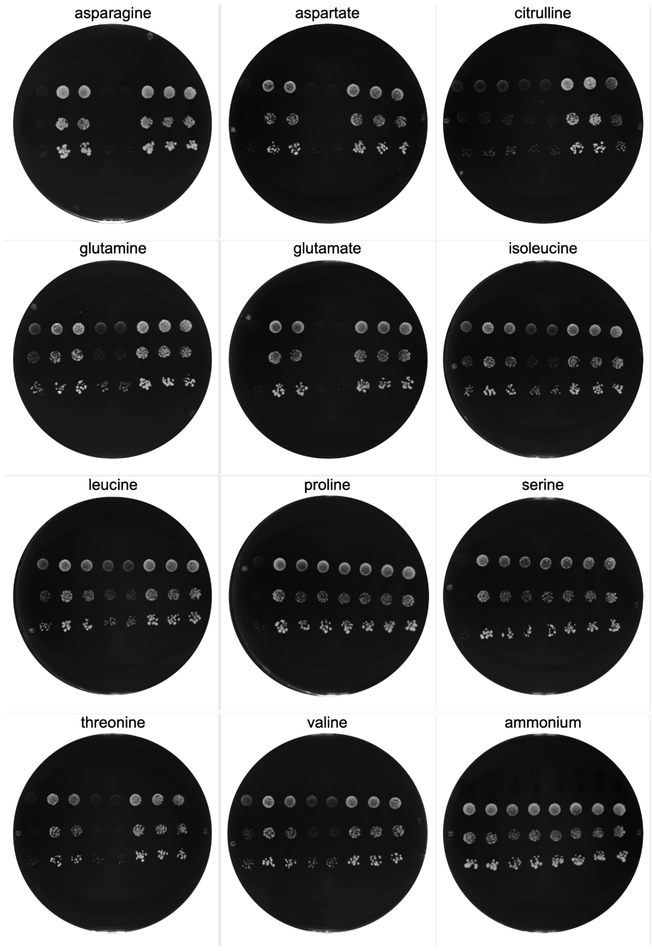


**Figure 2–figure supplement 2. *In vivo* evolution of *PUT4*.** Growth assays of Δ10AA pADHXC3GH-*PUT4* variants isolated from Asp and Glu evolution, spotted on minimal agar with 1 mM of the respective amino acid as the sole nitrogen source. Dishes were imaged after 15 days of incubation at 30 °C. Order of *PUT4* variants in each dish from left to right: negative control (*AGP1*-Cit9), *PUT4*-Glu2, *PUT4*-Glu1, *PUT4*-wild type, *PUT4*-wild type, *PUT4*-Asp3, *PUT4*-Asp2, *PUT4*-Asp1. Spotted are 5 µL of $OD_{600}$ of 0.1, 0.01, and 0.001.

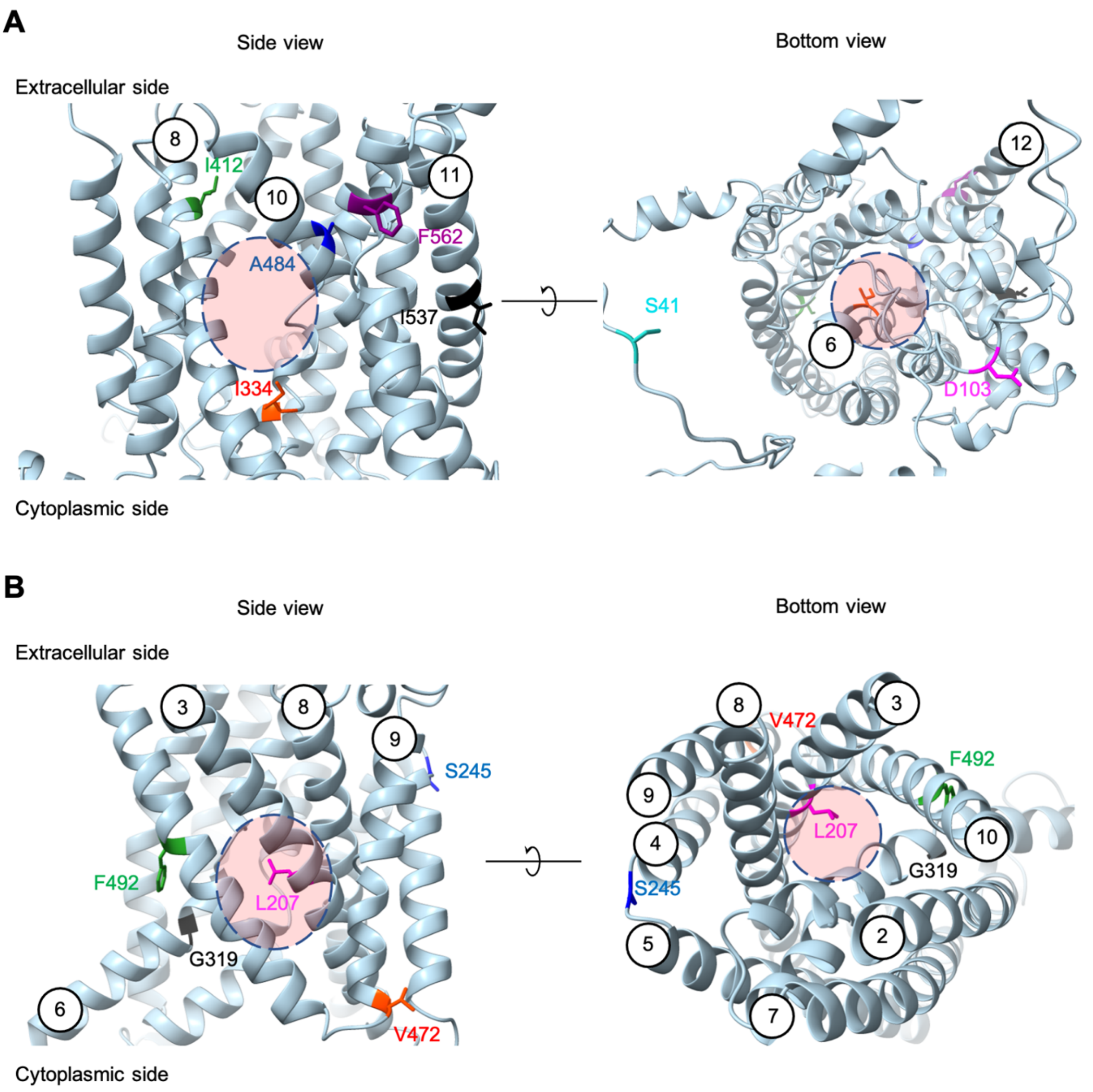


**Figure 2–figure supplement 3**. **Positions of the substituted amino acids found in the evolved mutants.** Side and bottom views of the AlphaFold models of *AGP1* (AF-P25376) (A) and *PUT4* (AF-P15380) (B) visualized in ChimeraX (1.3.0). The reported amino acids that were substituted in the evolved mutants are highlighted in different colors. The respective TMs are presented in circles. The predicted substrate binding site is represented as dashed circle.

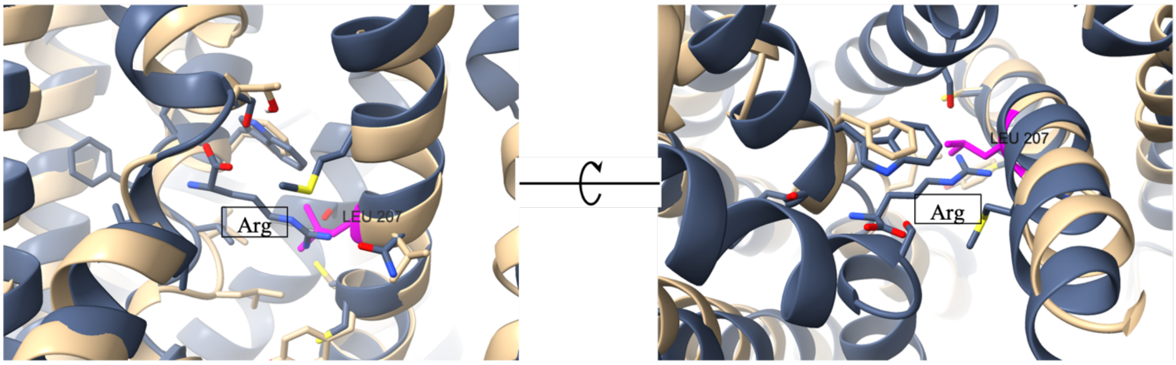


**Figure 2–figure supplement 4**. **Position of the L207S on the transporter's binding site.** Overlay of *AdiC* crystal structure (PDB: 3OB6; grey) and *PUT4* AlphaFold model (AF-P15380; beige). The original substrate of *AdiC* is present in the middle of the images ("Arg"; grey). The amino acids contributing to the substrate binding site are shown in sticks. The L207 of *PUT4* is shown in magenta, positioned in the binding site area. According to VAST Search (NCBI), the two structures share a 16.5% sequence identity in the superimposed protein parts, a structural similarity score of 22.15 and RMSD of 3.51 Å.

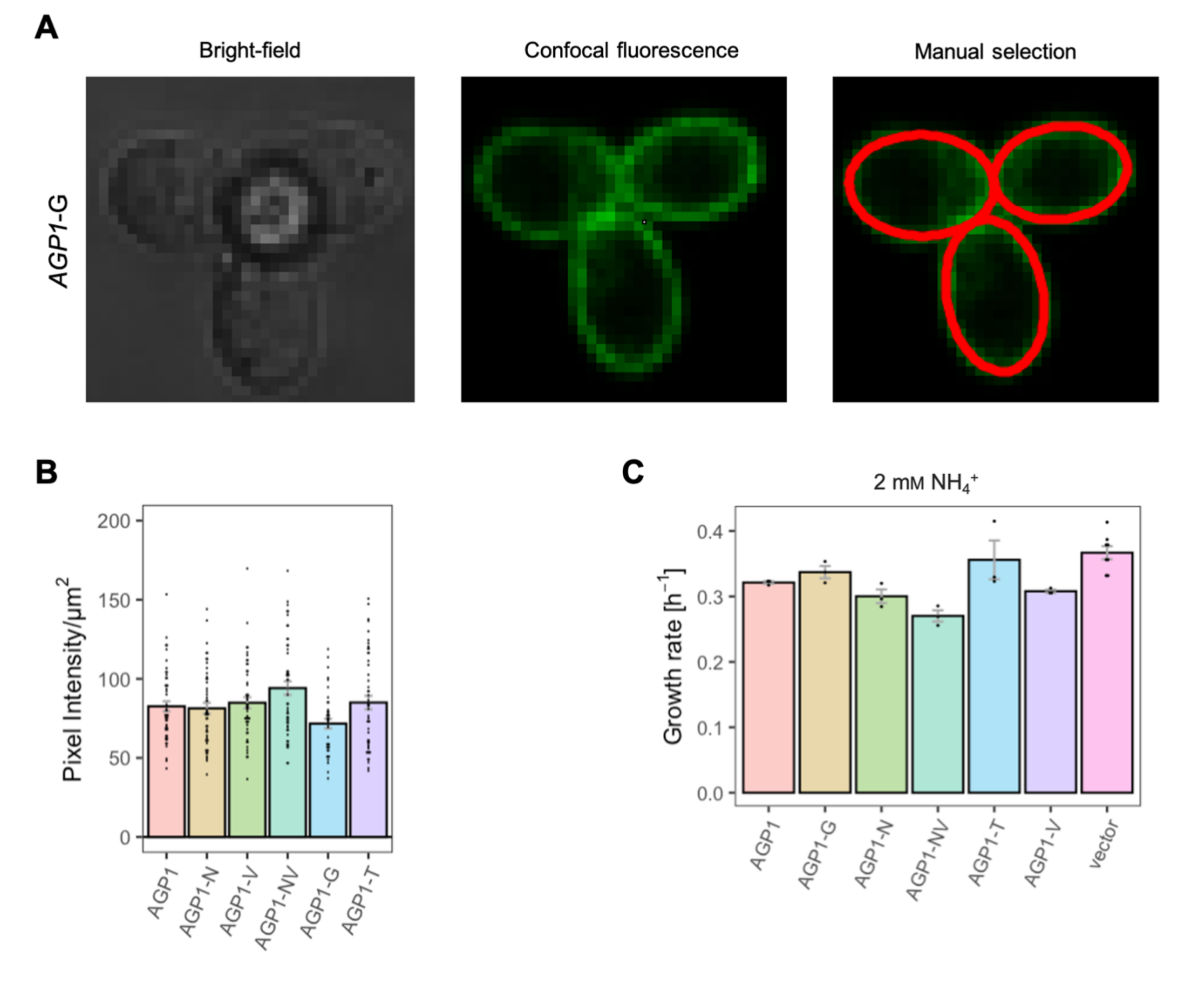


**Figure 3–figure supplement 1. Effects of evolved *AGP1* mutations on the surface expression and growth on non-amino acid nitrogen source.** (A) Localization of the *AGP1*-G variant in whole cells by fluorescence microscopy. The same cells are presented under the bright-field (left) and confocal fluorescence (middle) channels. The manual selection of the periphery of the cells (right) was performed in Fiji with 1 pixel width. (B) Surface expression of the *AGP1* variants. Error bars represent the SEM (n = 40-50). (C) Growth rate of the *AGP1* variants and the vector on 2 mM $NH_4^+$. Error bars represent the SEM ($n \geq 3$). One-way ANOVA with a Dunett's test showed no significant difference between the *AGP1* variants and wild-type for both (B) and (C).

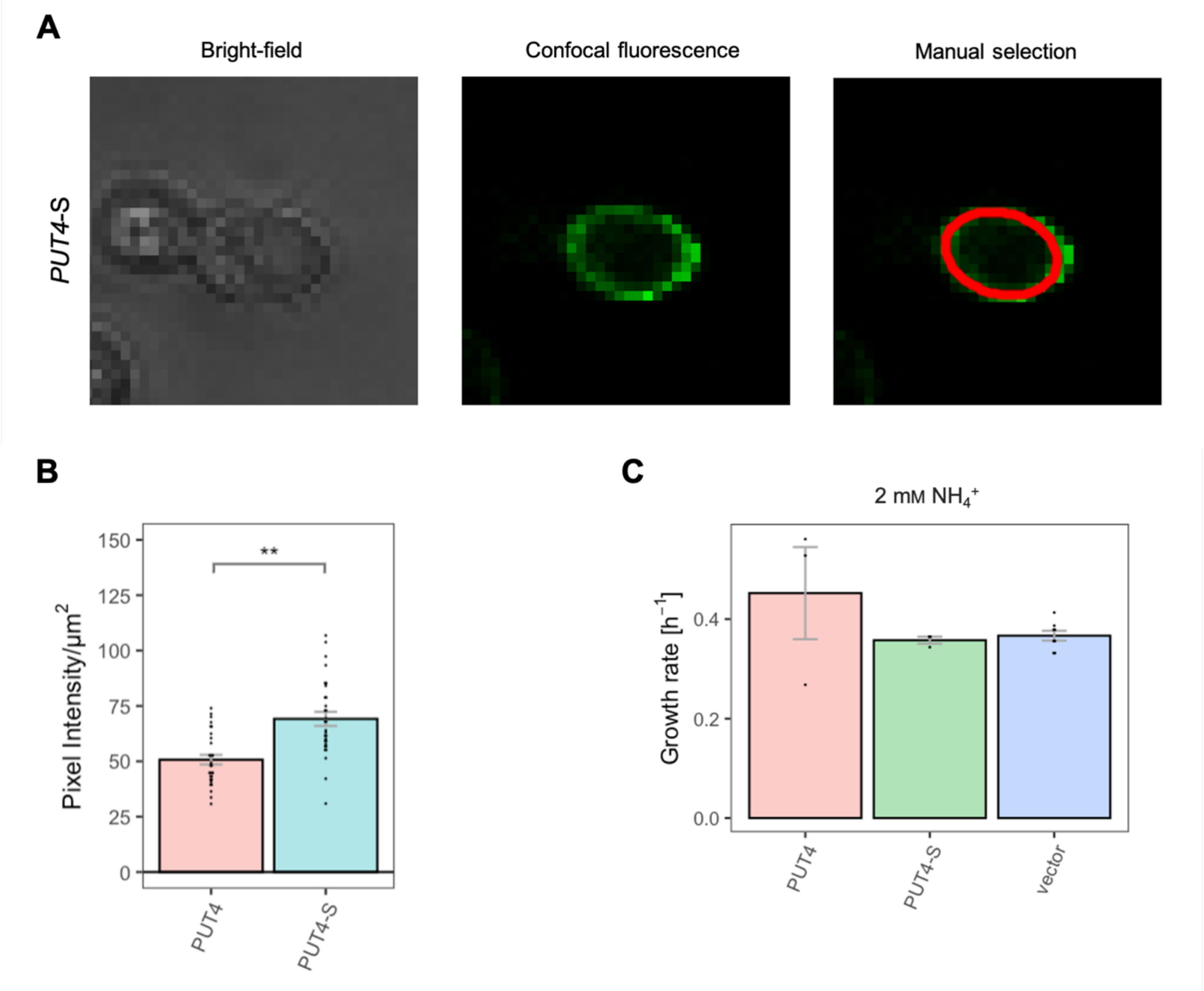


**Figure 3–figure supplement 2. Effects of evolved *PUT4*-S mutation on the surface expression and growth on non-amino acid nitrogen source.** (A) Localization of the *PUT4*-S variant in whole cells by fluorescence microscopy. The same cell is presented under the bright-field (left) and confocal fluorescence (middle) channels. The manual selection of the periphery of the cell (right) was performed in Fiji with 1 pixel width. (B) Surface expression of the *PUT4* variants. Error bars represent the SEM (n = 30). Asterisks indicate the degree of significant difference in pairwise comparison between the transporter-expressing variants (Student's t-test; ** $p < 0.01$, * $p < 0.05$). (C) Growth rate of the *PUT4* variants and the vector on 2 mM $NH_4^+$. Error bars represent the SEM ($n \geq 3$). Pairwise comparison (Student's t-test) showed no significant difference between the transporter-expressing variants.

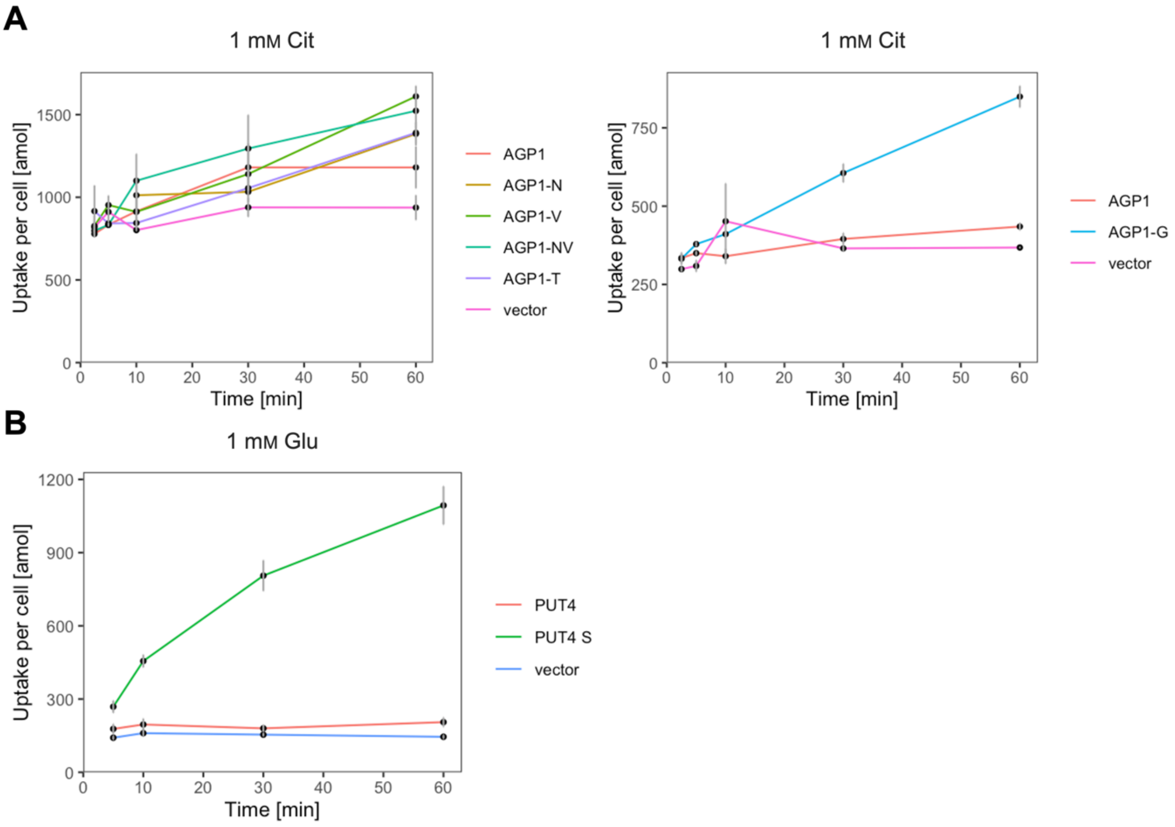


**Figure 3–figure supplement 3. The evolved variants support uptake of the respective amino acids.** (A) Uptake of 1 mM $^{14}$C-Cit by whole cells expressing different *AGP1* variants or none (vector). Error bars represent the SD (n = 3). Each graph represents an independent experiment. (B) Uptake of 1 mM $^{14}$C-Glu by whole cells expressing different *PUT4* variants or none (vector). Error bars represent the SD (n = 3).

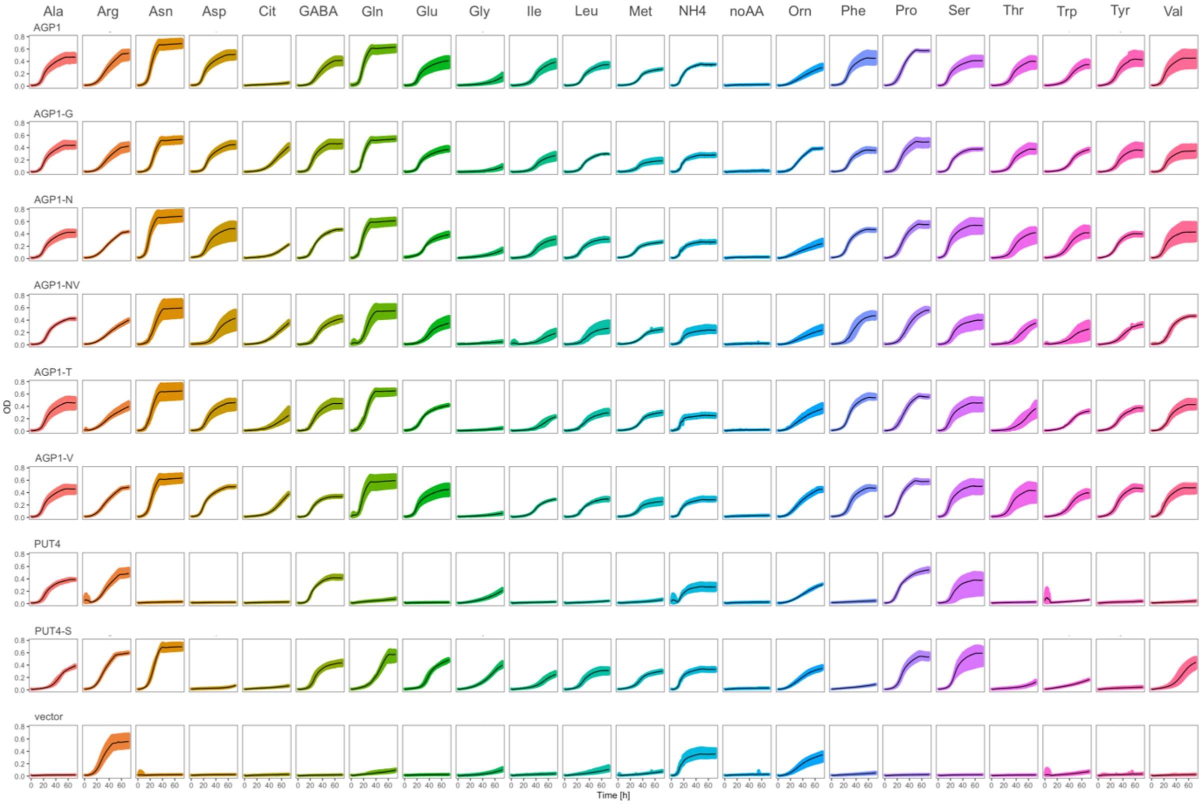


**Figure 4–figure supplement 1**. **The evolved variants affect the strain's growth on different amino acids.** Growth curves of Δ10AA expressing either one of the *AGP1* and *PUT4* variants from pADHXC3GH and the empty vector control on 2 mM of each amino acid. Black lines represent mean values of all measured curves (n ≥ 3). Colored areas represent the SD range.

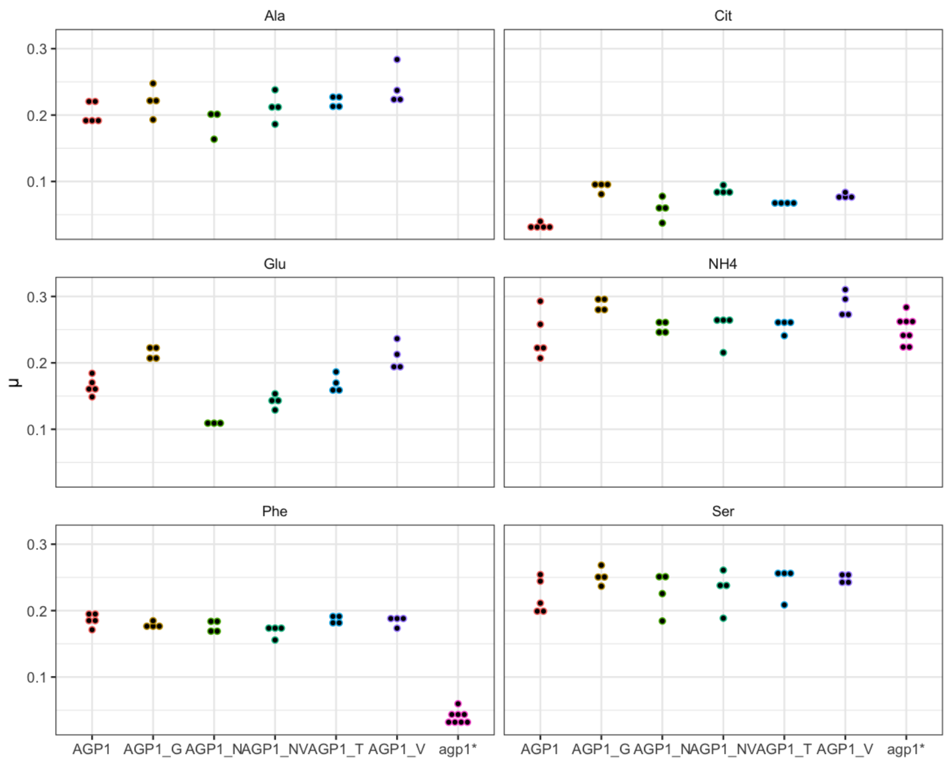


**Figure 4–figure supplement 2**. **Effects of evolved *AGP1* mutations on the growth rate on original substrates.** Repetition of the growth rate (μ) measurements of Δ10AA expressing either one of the *AGP1* variants from pADHXC3GH on selected substrates. Each point reflects the growth rate of one replicate culture. Note the slow growth of wild-type *AGP1* on Cit.

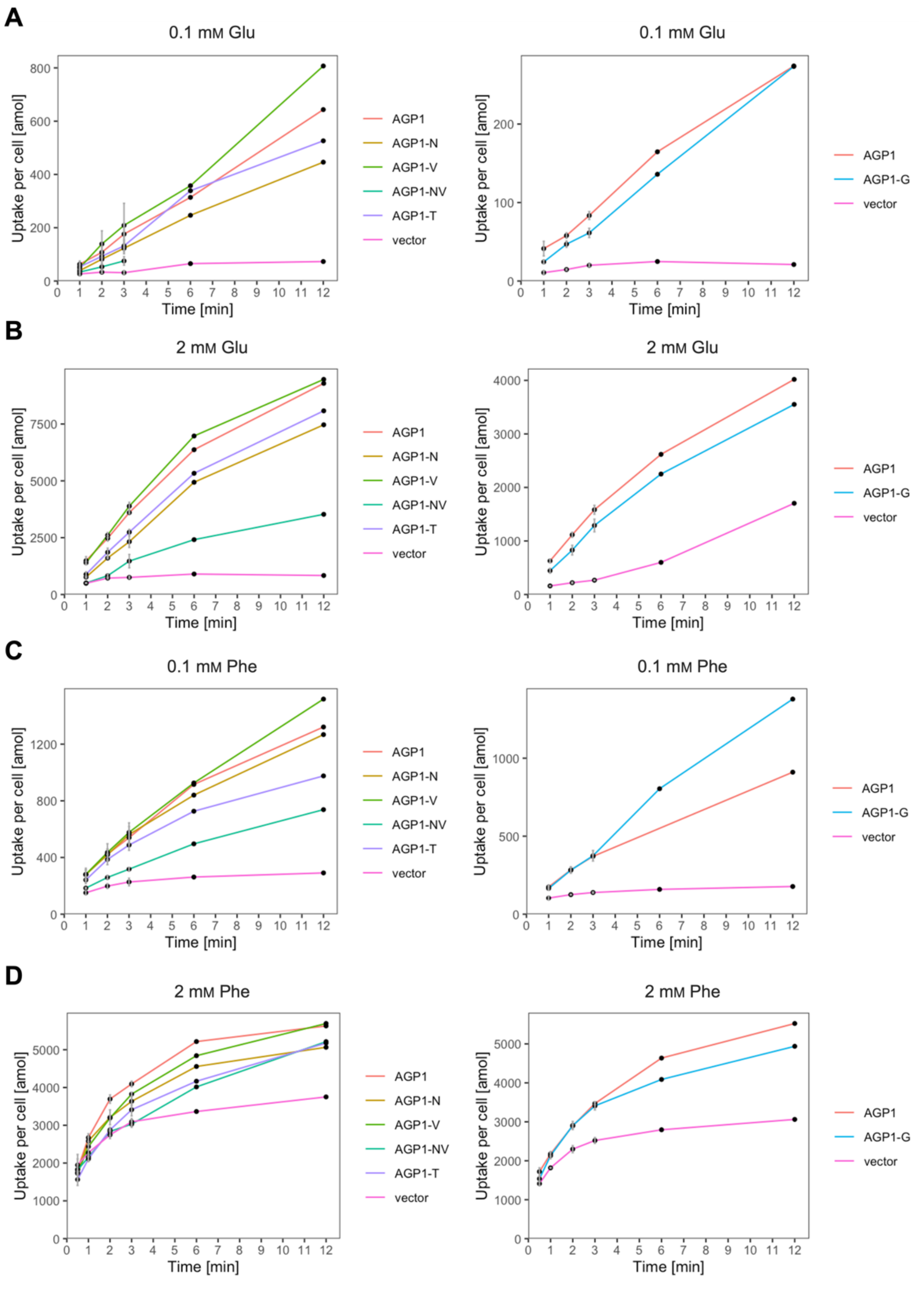


**Figure 5–figure supplement 1. The evolved *AGP1* variants support uptake of original substrates.** Uptake of 0.1 mM $^{14}$C-Glu (A), 2 mM $^{14}$C-Glu (B; left) or $^{3}$H-Glu (B; right), 0.1 mM $^{14}$C-Phe (C), 2 mM $^{14}$C-Phe (D) by whole cells expressing different *AGP1* variants or none (vector). Error bars represent the SD (n = 3). Each graph represents an independent experiment from the uptakes described in Figure 5.

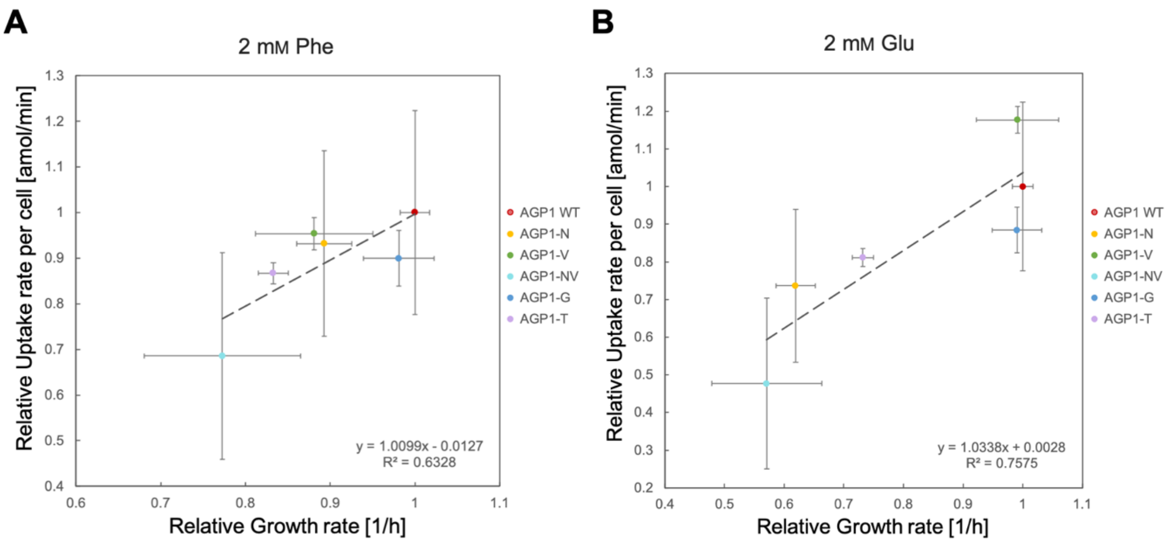


**Figure 5–figure supplement 2. Correlation between the fitness and uptake of original substrates for the evolved *AGP1* variants.** The uptake rate and growth rate on 2 mM Phe (A) or Glu (B) for the evolved *AGP1* variants is presented relative to that of cells expressing the wild-type protein. Error bars represent the SD ($n \geq 3$). The correlation is described by the linear regression trendline and the equation is shown in the graph.

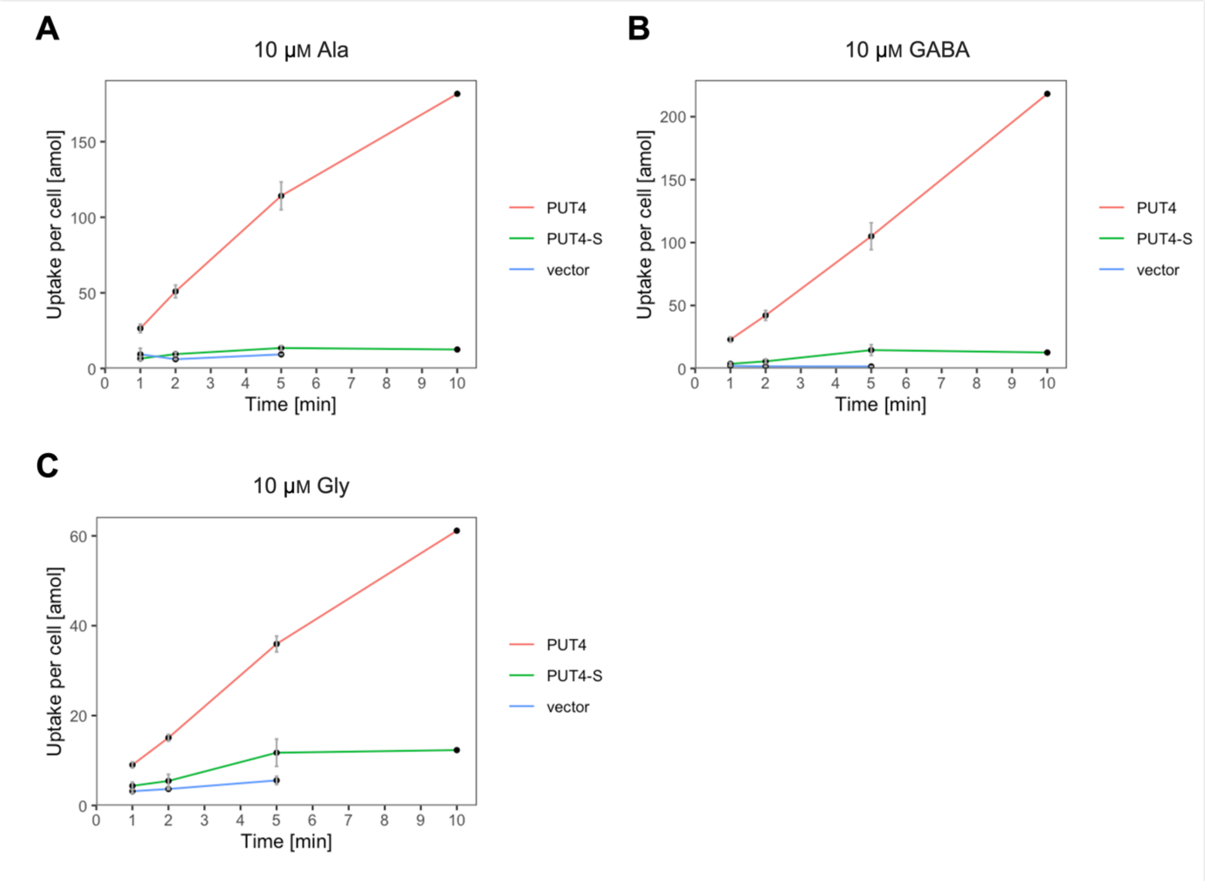


**Figure 5–figure supplement 3. The evolved *PUT4* variant supports uptake of original substrates.** Uptake of 10 µM $^{14}$C-Ala (A), GABA (B) or Gly (C) by whole cells expressing different *PUT4* variants or none (vector). Error bars represent the SD (n = 3). Each graph represents an independent experiment from the uptakes described in Figure 5.
